# ARID5B mutations cause a neurodevelopmental syndrome with neuroinflammation episodes

**DOI:** 10.64898/2026.01.15.698931

**Authors:** Hendrikus J. van Heesbeen, Nazim Rabouhi, Aurélie Gouronc, Angéline Preto, Justine Rousseau, Jade Charbonneau, Antonio Vitobello, Ange-Line Bruel, Anne-Sophie Denommé-Pichon, Estelle Colin, Bertrand Isidor, Sophie Nambot, Desiree DeMille, Pinar Bayrak-Toydemir, Nicola Longo, Boris Keren, Alexandra Afenjar, Jolien S. Klein Wassink-Ruiter, Ingrid P.C. Krapels, Hannah Titheradge, Gavin Ryan, Matias Wagner, Jill A. Rosenfeld, David R. Witt, Anirudh Saronwala, Yaping Yang, Annick Rein-Rothschild, Ortal Barel, Reena Jethva, Saskia B. Wortmann, Katharina Diepold, Kevin Rostasy, Lola K. Clarkson, Kathryn T. Drazba, Raymond J. Louie, Himanshu Goel, Outi Kuismin, Pekka Nokelainen, Jianling Ji, Ashley Mills, Matthew A. Deardorff, María Palomares-Bralo, María-Ángeles Gómez-Cano, Alberto Fernández-Jaén, Peter J. Hulick, Maureen Jacob, Benjamin Cogne, Kandamurugu Manickam, Xueqi Wang, Gail Graham, Bert Callewaert, Mercedes Zoeteman, Michael L. Raff, Marion Aubert Mucca, Médéric Jeanne, Grace Raines, Amy Crunk, Sureni V Mullegama, Taila Hartley, Kristin Kernohan, Kym Boycott, Philippe M. Campeau

## Abstract

Genetic disorders affecting the epigenetic machinery constitute a major group of neurodevelopmental conditions. Pathogenic variants in several ARID transcription factors—particularly *ARID1A*, *ARID1B*, and *ARID2*—cause Coffin–Siris syndromes, all characterized by intellectual disability (ID). These genes encode core subunits of the BRG1/BRM-associated factor (BAF) chromatin remodeling complex. In contrast, ARID family members that function in other regulatory complexes have remained largely unexplored in neurodevelopmental disease.

Here, we identify 29 individuals carrying heterozygous *ARID5B* variants, of which 24 (83%) introduce premature termination codons in the exceptionally long final exon, one affects the exon 9 splice donor site, and four are missense variants in conserved domains within the N-terminal half of the protein. Using a CRISPR–Cas9 knock-in mouse model harboring the p.Q522Ter variant, together with in vitro assays, we investigated the functional consequences of C-terminal *ARID5B* truncations.

All affected individuals presented with global developmental delay or ID—most commonly mild—and frequent speech and language impairment. Recurrent features included kidney malformations, behavioral difficulties, and recurrent infections of the respiratory and urinary tracts. Two individuals experienced central nervous system inflammation, and two infants presented with persistent pulmonary hypertension. Remarkably, 19 of 29 variants (66%) cluster within the first quarter of exon 10, are de novo, and escape nonsense-mediated mRNA decay (NMD), which we confirmed for two variants affecting seven individuals. Variants outside this region were inherited. Heterozygous mice exhibited developmental and behavioral abnormalities, while homozygous mutations was perinatally lethal. Truncations and a small deletion within a predicted nuclear localization signal (NLS) caused cytosolic mislocalization of ARID5B, whereas the isolated C-terminal half retained nuclear localization, suggesting an independent distal NLS.

Collectively, these findings define *ARID5B*-related neurodevelopmental disorder as a distinct clinical entity and reveal how disruption of specific *ARID5B* domains impacts protein localization, mammalian development, immune and neurobehavioral function.

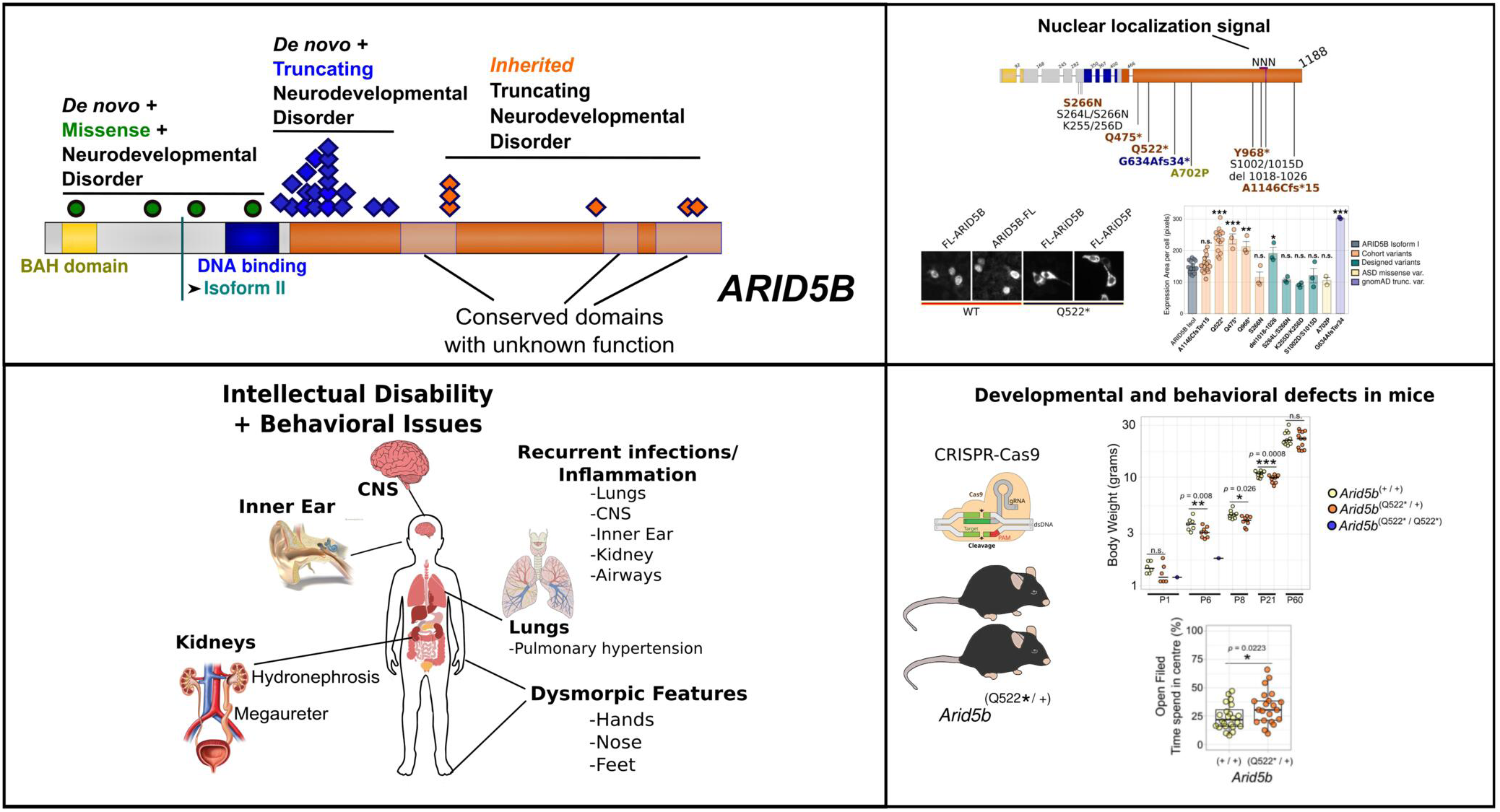

## Introduction

Epigenetic diseases that are caused by genetic variants have emerged as a major category of neurodevelopmental disorders (NDD). The AT-Rich Interaction Domain 5B (ARID5B [OMIM: 608538]), known as modulator recognition factor 2 (MRF2) or DESRT, belongs to the ARID transcription factor family which have key roles in gene regulation via chromatin remodeling and epigenetic modification^1^. Variants in several members of the ARID family have been associated with neurodevelopmental disorders, in particular *ARID1A* (Coffin-Siris Syndrome 2 (CSS2, [OMIM #614607]), *ARID1B* (Coffin-Siris Syndrome 1 (CSS1, [OMIM #135900])^2,3^ and *ARID2*, another member of the BRG1-associated complex (BAF) chromatin remodeling complex, leading to a Coffin-Siris-like syndrome^4^. ARID family members that are not members of the BAF complex have been less frequently associated with NDD. How-ever, via meta-analyses of a genomic study on intellectual disability (ID)^5^, we could associate a loss of function variant (LoF) in *ARID5B* with NDD. Notably, this variant would escape nonsense-mediated mRNA decay (NMD) as a consequence of the premature termination co-don (PTCs) being in the proximal part of the exceptionally long last exon, exon 10 of *AR-ID5B* (2.16 kb, 98th percentile for exon length based on NCBI human exon sizes)^6,7^.

Through gene matching, we initially identified three additional individuals with *de novo ARID5B* variants presenting with intellectual disability, speech and global developmental delay, learning difficulties and two with kidney defects. These individuals did not resemble the set of phenotypes typical of Coffin-Siris(-like) syndromes and lacked any disease-causing variants in known neurodevelopmental disorders. Moreover, all three truncating variants clustered in the first quarter of the long exon 10, similar to the reported variant, which prompted us to systematically characterize the clinical manifestations associated with *ARID5B* disruption.

Here, we present a cohort of 29 probands with developmental delay and rare ARID5B variants identified via the Matchmaker Exchange platform^8^. The individuals have divergent geographical and ethnic backgrounds and females and males are nearly equally distributed. Remarkably, the majority of individuals have truncating variants with premature codons in the long exon 10 that affect both TV1 and TV2 transcript variants while escaping NMD. The others four variants are missense variants in conserved missense intolerant domains in the N-terminal half of the protein.

## Materials And Methods

### Cohort selection criteria

Inclusion criteria were individuals with a predicted deleterious (premature termination codon or missense) variant in *ARID5B* and a neurodevelopmental disorder (NDD), in the absence of known NDD causing variants. The criteria for a NDD were either if patients presented with global developmental delay (all but one), or when speech was delayed, or when intellectual disability was diagnosed (ID). All the cases were isolated, except for three sisters, who inherited the variant from an affected mother. All variants were identified through exome or genome sequencing.

### Data and sample collections

Informed consent for genetic studies as well as the blood sampling and publication of photographs included here was obtained from parents or legal guardians. Written, signed forms were accompanied for consent to use the donated cells and images of the participating individuals or their parents. Lymphoblastoid cell lines (LCLs) were provided by Care4Rare Canada. Peripheral blood mononuclear cells (PBMCs) controls were generated following blood sampling and from cohort individuals provided by M.D. and R.J. and subsequently processed at the Centre hospitalier universitaire Sainte-Justine (CHUSJ) Institutional Mother-Child Biobank, following standard protocols.

### Mice

C57BL/6NJ mice were maintained in an established facility at the Centre de recherche Azrieli du CHUSJ (CRA-CHUSJ). Mouse husbandry and colony maintenance were performed according to the animal protocol approved (2022-3860) by the “Comité Institutionnel des Bonnes Pratiques Animales en Recherche” (CIBPAR). This committee is following the guidelines of the Canadian Council on Animal Care (CCAC).

### Generation of Arid5b*^emQ522*^* mice

The human variant NM_032199.3 c.1564C>T, p.Gln522Ter, was introduced into the mouse genome using CRISPR/Cas9. The mice were generated by the McGill Integrated Core for Animal Modeling (MICAM; McGill University, Montreal, Quebec, Canada). Briefly, the sgRNA (Synthego), Cas9 protein (IDT, catalog 1081058) and ssODN (ultramer, IDT) were microinjected into the pronucleus of C57BL/6N mouse zygotes with respective concentrations of 50:50:30 ng/μL. Embryos were subsequently implanted in CD-1 pseudopregnant surrogate mothers according to standard procedures approved by the McGill University Animal Care Committee (UACC). Oligonucleotides used were mArid5b-gRNA: 5’-AAGGCCAATGAAACTGACCA-3’ and ssODN:5’-TGGGAGCTGAATCTTTTTCAGGAGGCAAGGGAGGGCTTGGAAGCAGAGGGGCAA GGCCCTTATCTCCCATCTCCTCGGCCTCTTTCTCGCTGTTGGAA**gg**TT**aa**TCAGTTT CATTGGCCTTTTCTGGGTCTGCTCTGGACAC-3’. After weaning, the mice were transferred to the CRA-CHUSJ. Mice were backcrossed for at least 3 generations to C57BL/6NJ (Jackson). The presence of the variant was confirmed by Sanger sequencing. Mouse husbandry and experiments at CRA-CHUSJ were done according to the approved animal user 732-NAGANO protocol no. 2021-3228 by the coordinator of the CIBPAR, in accordance with the CCAC guidelines.

### Staining quantification

ImageJ software^9^ was used to quantify pictures of fluorescence semi-automated and blinded. For each data point in **Figure 5B**, 2 wells of a 12-wells plate were used per condition and for each well, 2 images were generated and quantified. For the quantification in **Figure 5B**, channels from the overlay ImageJ.tif files for DAPI and FLAG were split. The option Process>Binary>Make binary was used and ‘Analyze Particles’ to determine the region of interest (ROI). For the settings, a range of 20 to 1000000 pixels for FLAG and from 10 to 1000000 for DAPI was chosen and holes were included. This way, we verified that almost solely separated single nuclei were selected using the DAPI channel and FLAG staining images generated ROIs that corresponded to single cell expression, regardless of whether the staining was cytosolic and/or nuclear, this way selecting transfected cells. Next, the nuclear ROI was overlayed on the 16-bit FLAG-channel layer to quantify total FLAG staining per cell. Next, the DAPI ROI overlay was again used to fill up the nuclear ROI within the FLAG-16 bit image, masking the nuclear FLAG-staining but leaving the cytosolic expression. Then, we used the FLAG ROI again to quantify single cellular cytosolic staining. Nuclear staining per individual cell was determined as the difference between total FLAG-staining and cytosolic staining. Finally, each image output nuclear/cytosolic ratios for each cell were calculated per cell, generating several hundreds of measurements per image, then averaged per well and used for statistics and graphical presentation. This was repeated for three separate experiments and the averages of each experiment are plotted and used for statistical analyses. For the quantification in **Figure 5G**, one image per (n) per condition was taken and measurements were based on the size of the FLAG area per cell (imageJ, ROI per cell), see Rscript. Each data point is the mean of ROI surfaces of all individual cells in one coverslip/12-wells well. The wells were from three separate plates.

*See **supplemental information** for additional, more standardized experimental procedures and statistics*.

## Results

### *De novo* truncating *ARID5B* variants cause neurodevelopmental phenotypes

Assessing the potential deleterious effects of variants in the ARID family, other than those in the previously established critical neurodevelopmental genes *ARID1A* and *ARID1B*, we initially were compelled by a loss of function (LoF) variants in *ARID5B* in a genomic study on individuals with intellectual disabilities^5^. We next collected information on three additional individuals with developmental delay (DD) and *ARID5B* variants (not selecting for a specific type of variant) with exome or whole genome sequencing. Remarkably, all four variants caused premature termination codons (PTCs) at the beginning of the long final exon 10, leading to the truncation of at least half of the protein while predicted to escape nonsense-mediated mRNA decay. This prompted our hypothesis that truncating variants at the beginning of exon 10 can cause neurodevelopmental perturbations and motivated us to collect a larger cohort of *de novo ARID5B* variants, for which we also aimed to collect a more complete clinical profile.

With the help of Matchmaker Exchange platform tools^8,10^ we have assembled a cohort of in total 29 probands with neurodevelopmental phenotypes and *ARID5B* variants of which 24 (83%) are loss of function variants that have PTCs in exon 10 and one that affect the exon 9/10 splice site. Four others are missense variants in conserved protein domains in the N-terminal half of the protein. (**Fig. 3B,C**). Nineteen (68%) variants lead to gained stops or frame shifts in the first quarter of the long last exon 10, all of those that were assessed (14 individuals) were *de novo* variants. However, five out of six exon 10 truncating variants that were more dispersed over the other three quarters of exon 10 were inherited from three symptomatic mothers (including three sisters) and one non-symptomatic mother. Inheritance of a sixth C-terminal truncating variant could not be assessed.

A detailed summary of overlapping dysmorphisms and images can be found in **Figure 1**. In **Figure 2** a detailed summary of frequencies of other overlapping phenotypes can be found. Both the number of individuals and the percentages are noted, the latter are calculated by excluding individuals of whom a particular phenotype was not assessed at the time of writing. Detailed per patient information including phenotypes that occur only once can be found in **Supplementary table 1**.

**Figure 1.**
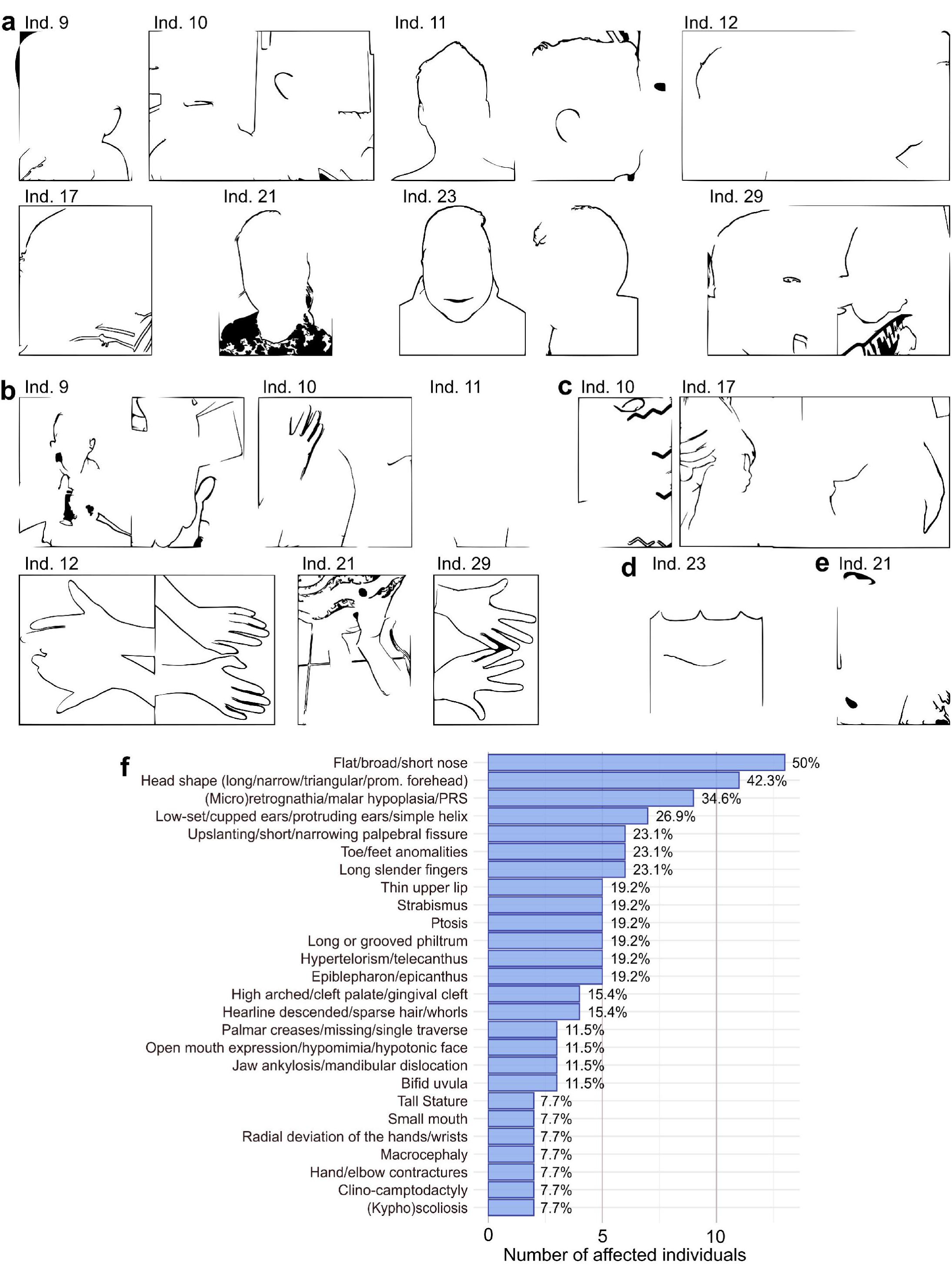
Dysmorphic features. (**A**) **Facial dysmorphic features**. **Individual 9**, with variant c.1379delA, p.Lys460SerfsTer19, has a broad nasal tip, protruding cupped ears, gingival cleft, micrognathia. **Individual 10**, with c.1398+1del, affecting the exon 9-10 splice site, has a prominent forehead, deeply groved philtrum, broad nose, and protuberant ear helices. **Individual 11**, with variant c.1419del, p.Glu474AsnfsTer5, has a high forehead, hypertelorism, telecanthus, high arched palate, ptosis, small mouth, long philtrum, malar hypoplasia, and hypomimia. **Individual 12**, with variant c.1420del, p.Glu474AsnfsTer5, has left ptosis, hypertelorism, long philtrum, a broad nasal tip, micro- and retrognathia. **Individual 17**, with variant c.1489dupA, p.Ile497AsnfsTer31, has mild ptosis, frontal bossing, telecanthus, a short nasal bridge, long philtrum. **Individual 21** (c.1587_1588del, p.Ala530ArgfsTer38) has a wide forehead and a broad nasal ridge. **Individual 23** (c.1804C>T, p.Gln602Ter) has almond-shaped eyes, slightly upslanting palpebral fissures, a hyperteloric appearance to eyes with telecanthus, and a bulbous nose with a slightly bifid nasal tip. **Individual 29** has a C-terminal variant (c.3435dupT, p.Ala1146CysfsTer15) inherited from his mother (not shown here), who both having a triangular-shaped face with ptosis, hypertelorism or telecanthus and a thin upper lip. (**B**) **Hand and foot features. Individual 9** had bilateral mild radial deviation of wrists, bilateral mildly overlapping toes. **Individual 10** camptodactyly of distal interphalangeal (DIP) joints, small hypothenar eminence contour. **Individual 11,** presented with the proximal phalanx of his fifth finger short and the base lower, as indicated by the position of the proximal interphalangeal crease. **Individual 12** presented with clinodactyly of his fifth finger, and his palmar creases are hardly visible, small hypothenar eminence contour. **Individual 21** presented long flat feet and increased sandal gaps. **Individual 29** presented with the position of the base of the small finger and the proximal interphalangeal joint are aberrant with a deviation of the hands with slender fingers, as seen for other individuals. (**C**) **Individual 10** presented with circumferential skin folds on upper (and lower) extremities, and prominent umbilicus (not shown). **Individual 17** presented with mild laxity of wrists, elbows, knees, and fingers, and out-toeing of his feet (suspect vertical talus) with folded skin and an umbilical hernia. (**D**) **Individual 21** presented with abnormal teeth and dentition. (**E**) **Individual 23** presented a lobulated, slightly bifid uvula. His congenital pes planovalgus of both feet, right single palmar crease, and narrowing bilateral distal palms are not shown. (**F**) **Overall frequencies** of recurring (more than one) dysmorphisms, with calculated percentages that indicate the frequencies as a percentage of addressed individuals. Only explicitly known features that were not adressed per indivdual are subtracted, for example when individuals are too young to asses certain phenotypes, or when complete assesment was not yet performed by the time of writing. Abbreviation: PRS = Pierre Robin Sequence.

**Figure 2.**
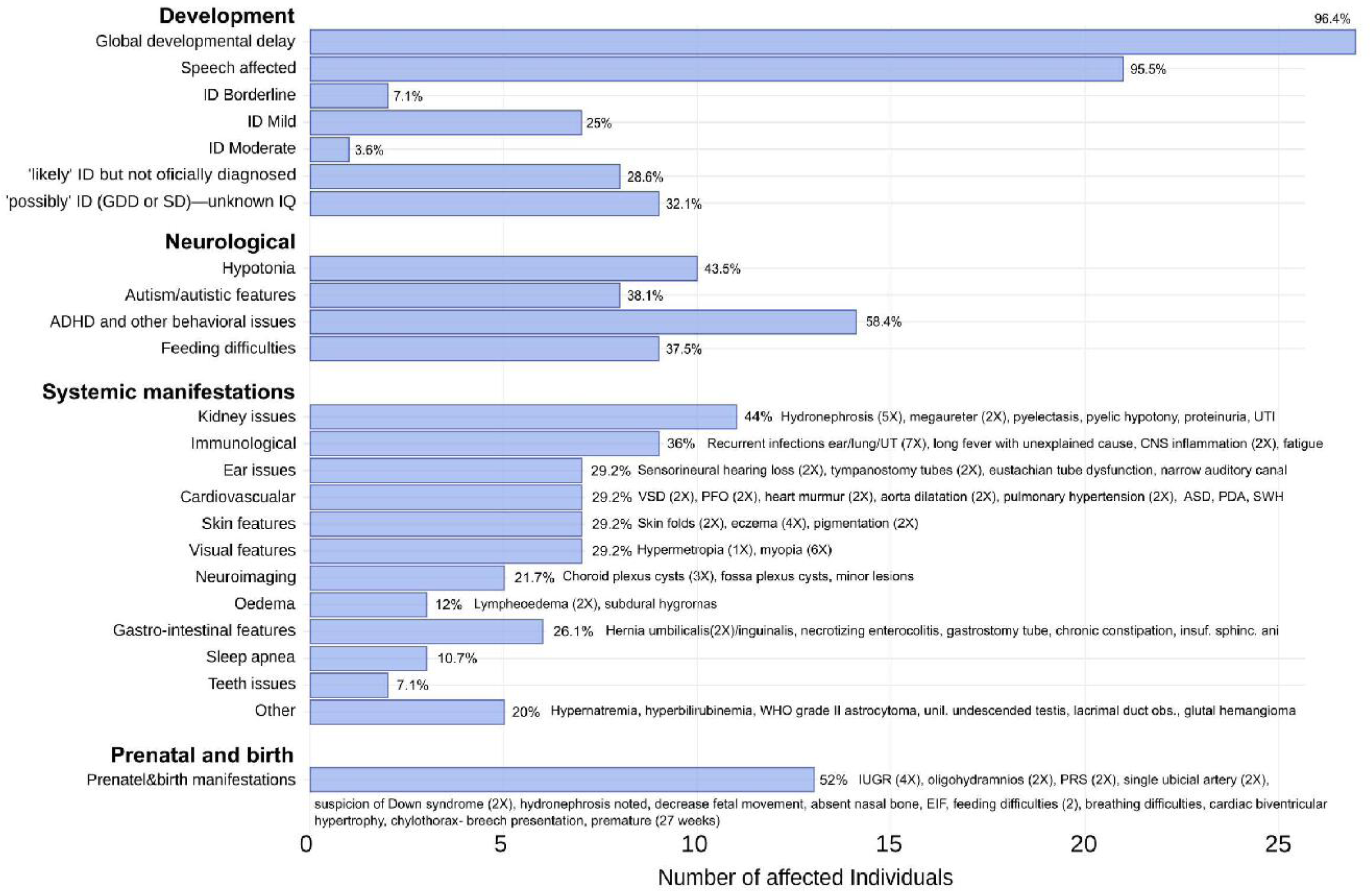
Phenotype frequencies. Frequencies of recurring phenotype features, except dysmorphism features, with calculated percentages that indicate the frequencies as a percentage of addressed individuals. Only explicitly known features that were not addressed per individual are subtracted, for example when individuals are too young to asses certain phenotypes, or when complete phenotype assessment was not (yet) performed by the time of writing. General categories are used to calculated frequencies of related features. Specific features with their recurrence (between parentheses) are written out next to the calculated percentage of each bar. **Abbreviations:** ID = Intellectual disability [OMIM: 156200], GGD = Global developmental delay, SD = Speech disorder/delay, ADHD = Attention Deficit&Hyperactivity Disorder, IQ = Intelligence Quotient, UTI = Urinary Tract Infection, CNS = Central Nervous System, SWH = Septal Wall Hypertrophy, ASD = Atrial Septal Defect, PDA=Patent Ductus Arteriosus [OMIM: 607411], VSD=Ventral Septal Defect [OMIM: 614431], PFO=Patent Foramen Ovale, WHO = World Health Organization, IUGR = Intrauterine Growth Restriction, EIF = Echogenic Intracardiac Focus, PRS = Pierre Robin Sequence.

Of the 29 probands, 14 females and 15 males with the age of their last follow up ranging from 0 to 29 years, all presented with a neurodevelopmental disorder. All individuals had global developmental delay (GDD), with the exception of one individual who was not assessed but suspected to have mild ID. Ten individuals were officially diagnosed with intellectual disability (ID [OMIM: 156200]), mostly mild (seven individuals), eight others likely have ID, but were not formally diagnosed at the time of writing, and all others were suspected to have cognitive challenges. Behavioral issues were frequent (18 individuals), including autism spectrum disorder (ASD [OMIM: 209850]) (8 individuals) and other behavioral issues (14 individuals), particularly attention-deficit/hyperactivity disorder (ADHD [OMIM: 143465]). Speech was affected in 21 individuals, including one individual with regressive language externalization leading to speech apraxia.

Frequent systemic manifestations that characterize the disease are kidney abnormalities (11 individuals) including hydronephrosis (five individuals), megaureter (two individuals), pyelectasis and pyelic hypotony. Hypotonia (10 individuals) and feeding difficulties (nine individuals) were common, with three individuals presenting with oropharyngeal dysphagia. Visual impairments (seven individuals), including myopia (OMIM: 160700) (6 individuals) and in addition three individuals with strabismus (OMIM: 185100). Ear/auditory issues (seven individuals) included sensorineural hearing loss (2 individuals) and tympanostomy tube placement (two individuals).

Immune dysfunction was noted in nine individuals, mostly with recurrent infections (seven individuals), two individuals presented with rare forms of CNS inflammation and another individual was hospitalized for a prolonged fever with unknown cause. Individual 27 (8-year-old girl) developed autoimmune cerebellitis and was unresponsive to therapy, while individual 23 (7-year-old boy) had recurrent acute disseminated encephalomyelitis (ADEM), but responded to therapy. Both were treated with corticosteroids and intravenous immunoglobulin (IVIG) and experienced seizures related to neuroinflammation. Two infants presented with persistent pulmonary hypertension, which was the predominant cause of death in individual 19 at six months of age. Finally, out of the four individuals that had missense variants, two presented with macrocephaly and another one with (mild) microcephaly with wide ventricles and corpus callosum atrophy that appeared after the age of 10 months. Macrocephaly was not reported for truncating variants, while individual 15, a two-year-old girl with a truncating variant, presented with a head circumference <0.4th centile. Finally, intrauterine growth retardation was reported 4 times, while post-natal growth was normal, with two individuals having a tall stature, and most individuals with truncations at the beginning of exon 10 being on the tall side.

A very common craniofacial feature was the shape of the nose, generally shorter and broader, a bit bulbous, often with a lower nasal bridge (13 individuals) (**Figure 1A,B**). 10 individuals had an uncommon head shape, with a more prominent forehead and narrow face, frequently with micro/retrognathia (8 individuals). Five individuals presented with hypertelorism, ptosis and, nine individuals presented with thin upper lip, long philtrum, or narrow mouth. Skeletal abnormalities included toe/foot anomalies (7 individuals), radially deviated wrists (2 individuals), long and slender fingers or clino-comptodactyly (6 individuals) and abnormal palmar creases (6 individuals).

### Premature termination codons cluster in the first quarter of exon 10 and escape NMD

Nineteen out of 29 variants (66%) cluster in exon 9 (only frameshift) and early exon 10 (frameshift and nonsense), with all PTCs being in the first quarter of exon 10 and one variant that affects the exon 9 / exon 10 splice donor site (**Fig. 3B**). Since PTCs in such exceptionally long exon 10 (2.16 kb, >95th percentile for length)^6^ are known to escape NMD, we quantified and sequenced the exon 9 – exon 10 border at the mRNA level to confirm the absence or strongly reduction of NMD in two lymphoblast cell lines (LCLs) and primary peripheral blood mononuclear (PBMCs) of two patients generated from blood collected at different ages. The variant affecting the donor of the LCLs was five times recurring in our cohort (p.Ile497AsnfsTer31). We indeed found no signs of NMD (**Fig. 3E, F**). Unlike our cohort, gnomAD LoF variants are dispersed, with only 3/23 (13%) clustering in the first quarter of exon 10, compared to 19/24 (79%) in our cohort (**Fig. 3B**)^11^. Moreover, all tested probands (5 out of 6, of which three sisters) were inherited, suggesting that the approximate location of the PTC could be relevant for the phenotypic outcomes. Less truncated, longer ARID5B protein could lead to milder phenotypes if they contain additional translated C-terminal domains that are still functional. One particular protein domain between amino acid 610-650, that is intolerant to missense variants, is lost in all truncations that cluster at the beginning of exon 10 (Fig 3B,C) but still encoded in all inherited truncations that affect the latter three quarters of exon 10. Moreover, the regional clustering of pathogenic LoF variants mirrors that seen in other epigenetic regulators (e.g., *KAT6B* [OMIM: 612990], *ASXL1* [OMIM: 612990], *ASXL3* [OMIM: 615115]).

**Figure 3.**
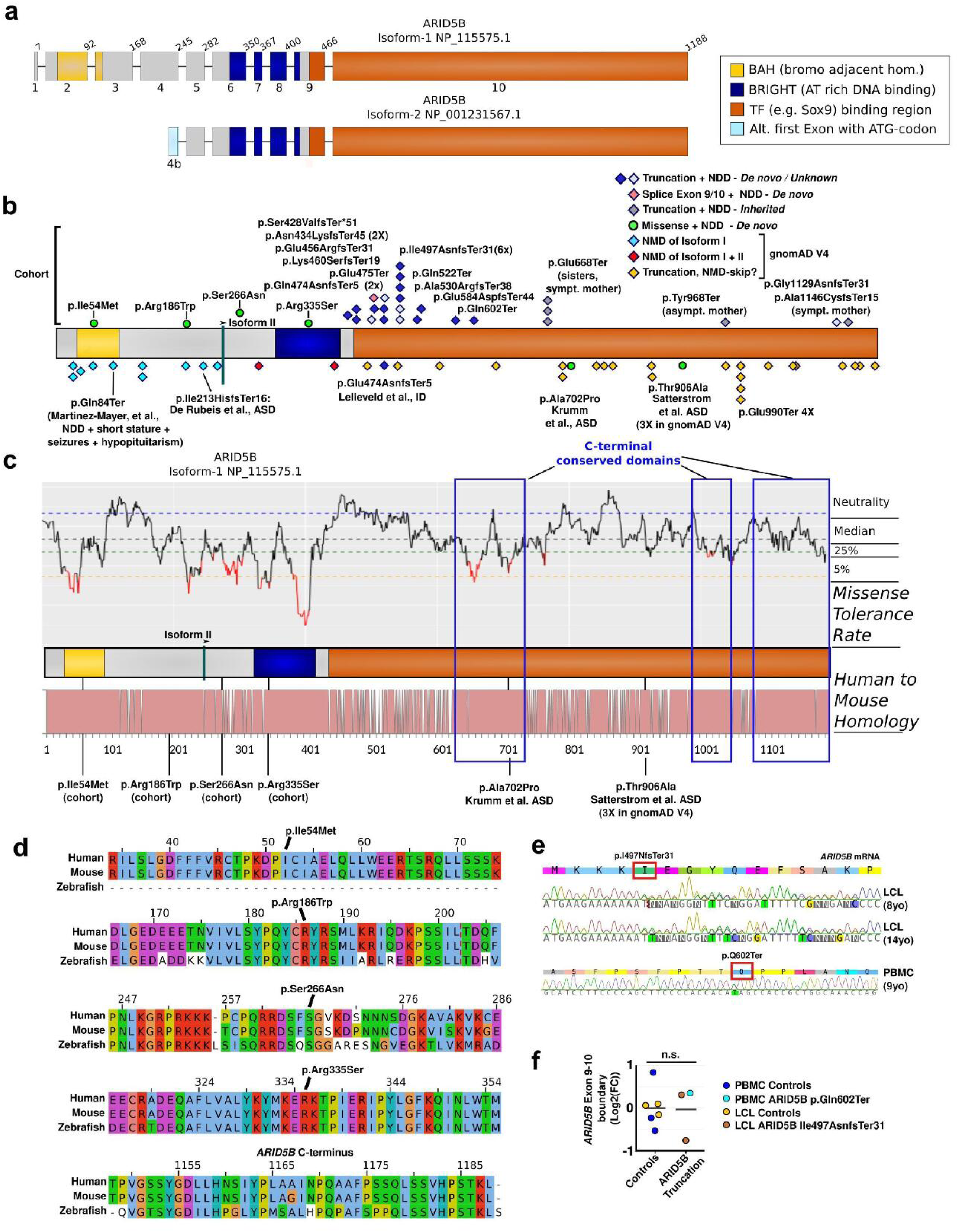
Schematic representation of *ARID5B* variant types and their consequences. (**A**) The two main isoforms generated from the curated transcript variants 1 and 2. The shorter isoform uses an alternative exon, here named 4b (light blue). Further, the two known domains, the BAH and the DNA binding BRIGHT domain are shown in yellow and dark blue, respectively. Sox9 has been shown to associate with the latter 2/3 of the protein (shown in orange), downstream of the BRIGHT domain. Variant p.(Asn434LysfsTer45) in this cohort has been included in the OMIXCARE cohort as well^39^. (**B**) Linear representation of isoform I, with the locations of the variants described in the current cohort on top and variants retrieved from gnomAD V4 or genomic studies below the protein bar. Diamonds indicate *truncating* variants. Light blue diamonds are predicted to cause NMD of only the long transcript variant 1. Red diamonds indicate variants that are predicted to cause NMD of both transcript variants. Dark blue variants, mainly found in the cohort, are both proven and predicted truncating variants associated with ID. The yellow diamonds represent truncating variants predicted to skip NMD, but with unknown phenotypes (retrieved from gnomAD V4). Green dots represent *missense* variants associated with neurological perturbations (four from the cohort, two from ASD studies^2,14^. (**C**) Isoform-I is represented with on top aligned the missense tolerance rate (MTR). The MTR indicates the running average *P*-value of synonymous versus observed missense variant. A lower value indicates a lower missense tolerance^40^. At the bottom, the homology between mouse and human ARID5B is represented with each non-conserved amino acid represented as a gap. Critical regions like the BRIGHT domain show a low missense tolerance and high conservation. Three mouse-to-human conserved domains are indicated with blue boxes. (**D**) Zebrafish-to-mouse-to-Human conservation of the loci with cohort missense variants (4X) and the ARID5B C-terminus. In (**E**), to confirm the expression of stable mutant RNA transcripts (escaping NMD), Sanger sequencing was performed on cDNA generated from RNA purified from patient cell lines, following DNAse treatment, and using primers that generate an amplicon that crosses the exon 9-10 junction (further avoiding amplification of DNA instead of RNA). The expression of mutated RNA with similar Sanger sequencing peak depth as wild type RNA. In (**F**), we validated this by quantifying the RNA levels and comparing normalized exon 9/exon10 RNA levels between controls and cell lines from patients with ARID5B variants, observing no effect on average RNA levels. Three PBMC control cell lines (blue dots) and three LCL control cell lines (yellow dots) were compared with two LCL clones generated at two different ages from individual 14 (c.1489dupA, p.Ile497AsnfsTer31; brown dots), and individual 23 (c.1804C>T, p.Gln602Ter; light blue dot). The line indicates average expression.

The four missense variants in our cohort resided in highly conserved, missense-intolerant regions and were not found in gnomAD V4; p.Ile54Met (BAH domain, but unknown target), pArg186Trp (domain of unknown function), p.Ser266Asn (domain of unknown function) and p.Arg335Ser (BRIGHT/ARID, DNA-binding domain). Especially the latter is in a strongly missense intolerant domain, and loss of a positively charged arginine at the end of the first alpha-helix in the the ARID5B DNA-binding domainis likely to affect binding to the negatively charged DNA^12^. All were conserved from mouse to human, and all but p.Ile54Met from mouse to zebrafish (Fig. 3C,D). Notably, in gnomAD V4^11^, the p.Ser266Gly substitution can be found 12 times.

Eight individuals presented with ASD, with only four of those being truncating at the beginning of exon 10, while three out of four missense variants, among which those residing in the BAH and BRIGH domain, were associated with ASD. We also noted two ASD-associated missense variants (p.Ala702Pro, p.Thr906Ala) in genomic studies^13,14^. Eventhough p.Ala702Pro was absent in gnomAD, p.Thr906Ala was identified four times in gnomAD V4. Another ASD-associated variant (e.g., p.lIe213HisfsTer16) affected only transcript variant 1 (TV1) (**Fig. 3B**).

### *Arid5b^(Q522*/+)^* mice exhibit developmental and behavior phenotypes

To further establish deleterious developmental consequences of truncating *ARID5B* variants, we generated knock-in mice that are carriers of the truncating variant of individual 20, a gained stop at the beginning of exon 10. Following repeated backcrossing and Sanger sequencing validation of the mutation and the ends of the donor DNA arms (**Fig. 4A**), we first tested the viability of mice heterozygous and homozygous for the variant. We did not observe any effect on the one-year survival rate of heterozygous mice, but homozygous mice generally died shortly after birth (**Fig. 4B**). Of note, with manual supplementation of food, some mice were able to survive slightly longer. Since homozygous mice remained very small and vulnerable, we decided not to proceed with their characterization for ethical reasons.

**Figure 4.**
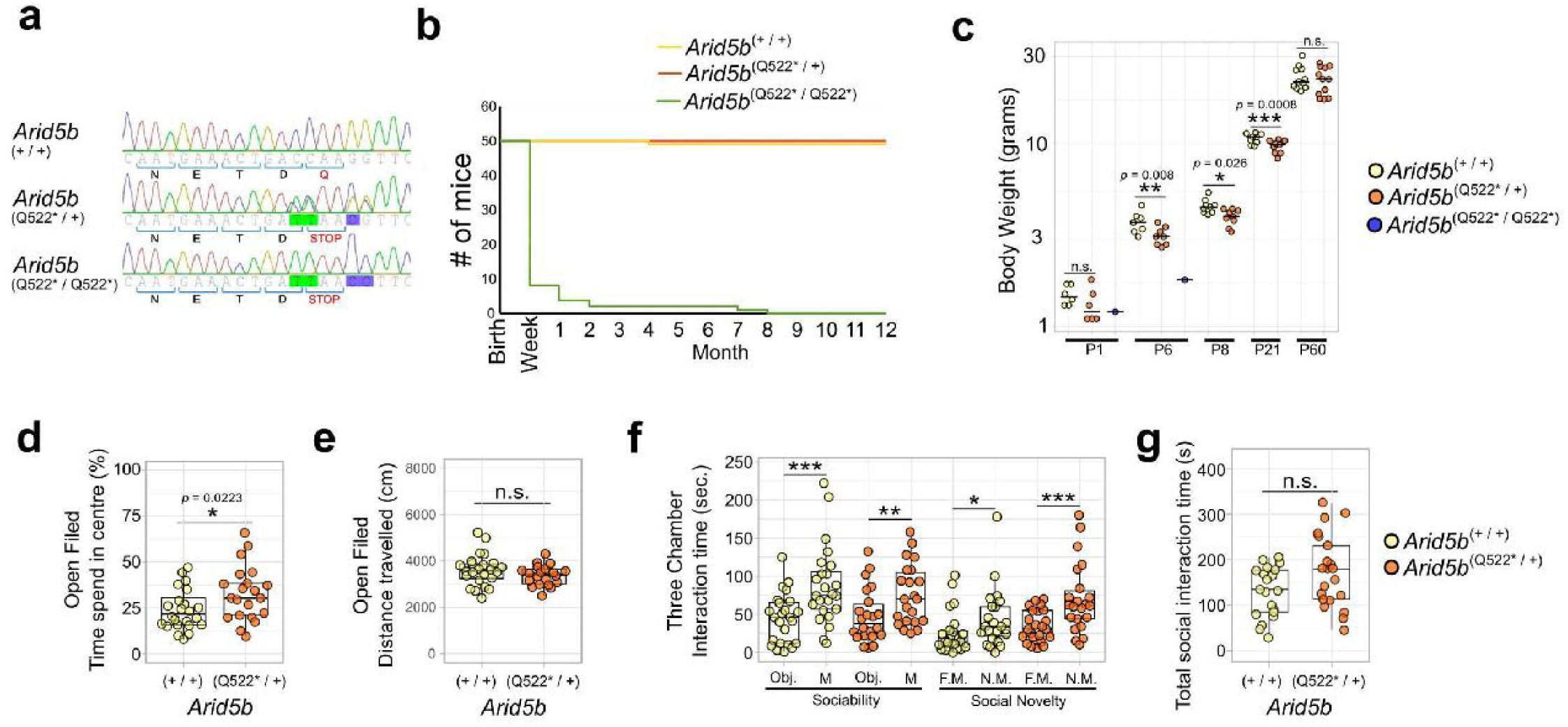
Mouse development and behavior are affected in *Arid5b^emQ522^*\* mice. (**A**) Sanger sequencing-based validation of genotype showing the four induced DNA mutations leading to a gained stop at Q522. (**B**) Kaplan-Meier survival plot over one year. The majority homozygous *Arid5b^Q522^*\* mice (green line) die in the first postnatal days. (**C**) Body weights of *Arid5b^Q522^*\* mice across development. Independent ages (P1, P6, P60) were analyzed using one-tailed t-tests to assess a reduction in body weight, comparing wild type with heterozygous mice. Repeated measurements at P8 and P21, obtained from the same cohort of mice, were analyzed using a linear mixed-effects model to account for repeated measures. Body weights at P60 were corrected for sex to account for naturally lighter female weights. Heterozygous *Arid5b*^Q522*^ mice have significantly reduced weight at P6 (*P* = 0.0084, df = 13), P8 (*P* = 0.0265, df = 22.10), and P21 (*P* = 0.00076, df = 22.10), but not at P60 (df = 21), P1 was inconclusive (*P* = 0.129, df = 10). (**D**) Open field test time spent in the center of the open field box as a percentage of total time spent in the open field box per mouse (wild type mice yellow dots/left box; *Arid5b*^Q522*^ mice orange dots/right box). A two-tailed t-test showed a significant increase in time spent in the center (*P* = 0.0223, df = 42). One outlier was removed. (**E**) The total distance covered during the total time spent in the open field box per mouse (wild type mice yellow dots/left box; *Arid5b*^Q522*^ mice orange dots/right box). A two-tailed t-test was inconclusive (*P* = 0.1852). (**F**) Three-chamber sociability test. Left side (“Sociability”): difference in time spent exploring an object versus a mouse for wild-type (yellow) and Arid5b*^Q522*^*mice (orange) mice. Right side (“Social Novelty”): difference in time spent exploring a familiar versus a novel mouse. Expected social behavior was confirmed using one-tailed t-tests to asses increased time spent with a mouse vs object or novel mouse vs familiar mouse. (**G**) Total social interaction time (summed across the sociability and social novelty experiments) in wild-type (WT, yellow) and *Arid5b*^Q522*^ (orange) mice. A one-tailed (*H*₁: reduced social interaction in heterozygous mice) t-test was not applicable, since the mean interaction time was increased in *Arid5b*^Q522*^ mice. A two-tailed t-test comparing total social interaction time between genotypes was inconclusive (WT = 132.17 ± 56.58 s, *n* = 21; Q522* = 173.64 ± 79.78 s, *n* = 21; *t*(36.1) = –1.94, *P* = 0.0598). Three outliers were identified and removed using Grubbs’ test — two in the wild-type group and one in the *Arid5b*^Q522*^ group. Statistical significance is indicated as follows: *P* < 0.05 (*), *P* < 0.01 (**), and p < 0.001 (***).

Even though heterozygous mice showed no signs of reduced one-year-viability, their lower weight throughout development suggested a developmental defect that seems to recover into adulthood (**Fig. 4C**). In addition, heterozygous mice showed different behavior in the open field test, being more frequently in the center than wild type mice (**Fig. 4D, E**). Similar patterns of behavior are seen in other mouse models of ID or ASD, like Fragile X syndrome (*Fmr1* disruption), Bosch-Boonstra-Schaaf Optic Atrophy Syndrome (*Nr2f1* disruption), *AUTS2*-related syndrome (*Auts2* disruption)^15–17^. However, we did not observe sociability or social novelty defects, as heterozygous mice showed a normal preference for a mouse over an object and for a novel mouse over a familiar mouse in the 3-chamber sociability test(**Fig. 4F**). Also the overall time that *Arid5b^(Q522*/+)^* mice spent on social exploration did not significantly change (**Fig. 4G**).

### Truncated ARID5B delocalizes into the cytoplasm

To investigate the protein consequences of truncating variants in exon 10, we constructed open reading frames (ORFs) encoding variously tagged ARID5B variants (see Methods). As expected, overexpression of ARID5B in HEK293T cells resulted in near exclusive nuclear localization under standard growth medium conditions, regardless of the tag added (**Fig. 5A**). However, introducing the nonsense variant of individual 20 (c.1489dupA, p.Gln522Ter) resulted in near complete delocalization of the truncated protein in the cytoplasm (**Fig. 5A**). Surface plots overlaying FLAG and DAPI staining further showed the strong extra-nuclear localization together with the nuclear absence of the p.Gln522Ter variant in more detail (**Fig. S2A**).

**Figure 5.**
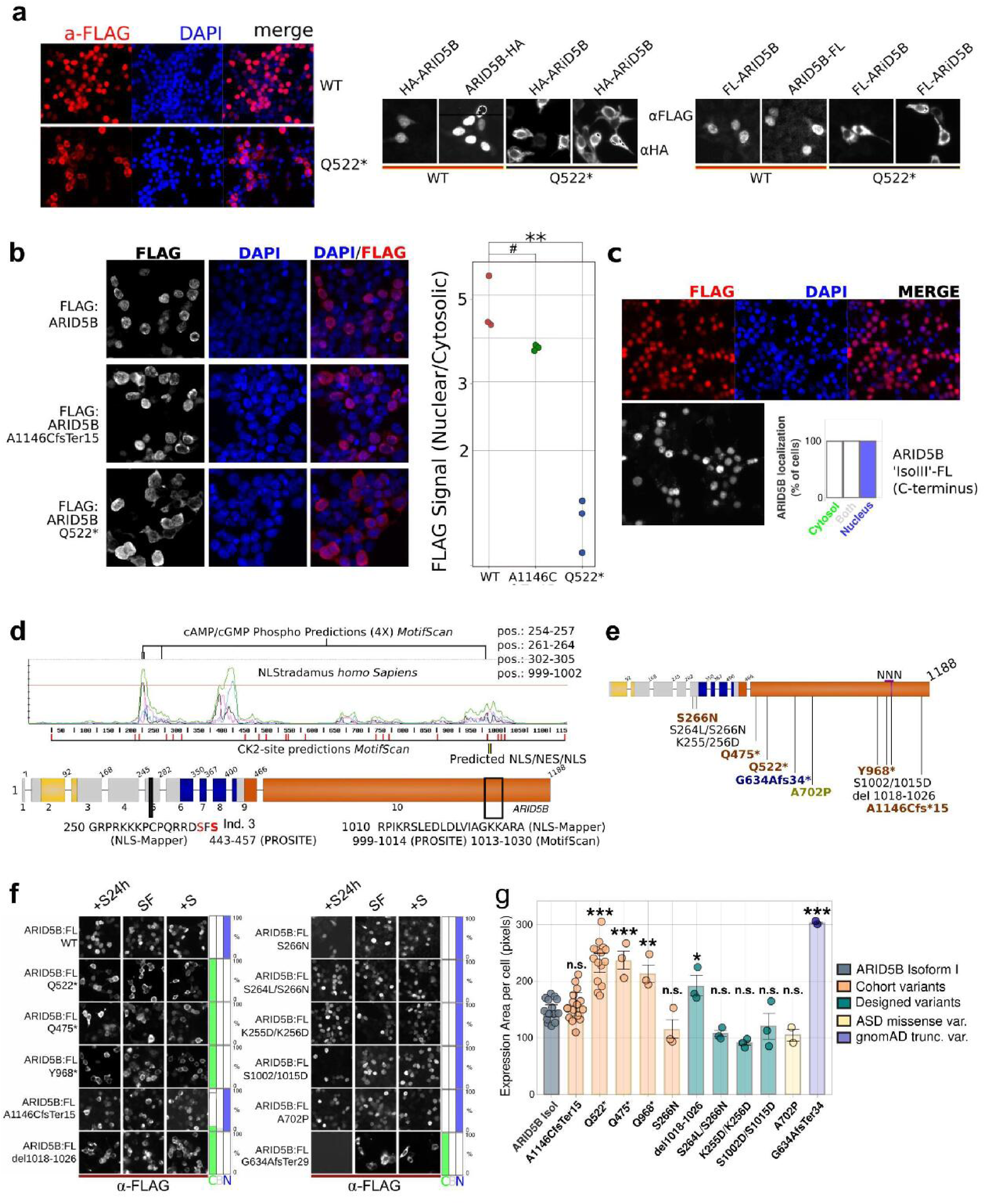
Terminating variants affect cellular localization with divergent effect sizes. (**A**) Overexpression with both C- and N-terminally, HA- or FLAG-tagged ARID5B isoforms in HEK293T cells show a strong preference for nuclear localization. The N-terminally HA or FLAG-tagged ARID5B, containing the variant of individual 10 (p.Glu522Ter), strongly locate in the cytosol instead. (**B**) Ratio of nuclear versus cytosolic ARID5B of (FLAG-tagged) wild-type, p.Glu522Ter, and the most C-terminal variant in our cohort detected for individual 22 and his mother (p.Ala1146CysfsTer15). The dots represent averages per experiment (see **Methods**). Two-tailed, two-sample t-tests (equal variance) compared each mutant to WT, with Bonferroni correction (α = 0.05). WT vs A1146C fsTer15: t(4) = 2.34, adjusted p = 0.176 (#), Cohen’s d ≈ 1.87 (ns). WT vs Q522*: t(4) = 7.39, adjusted p = 0.00412, Cohen’s d ≈ 6.49 (**). In (**C**), FLAG-tagged predicted isoform III cellular localization showing nuclear localization. (**D**) Analysis of ARID5B sequence using various structural prediction tools. The graph shows predicted nuclear localization sites (NLSs) generated by NLStradamus representing 3 Hidden Markov Modeled NLS states^18^. The elevated/peak signals predict three regions that could regulate nuclear localization. The most C-terminal elevated signals co-localize with predicted NLS and nuclear export signals (NES) predicted by other tools, here shown by name of the tool and amino acid sequence. Furthermore, the most abundant kinase signal transduction target sites that Motif Scan/Prosite^19^ predicted were Casein kinase 2 (CK2) sites, that are scattered and marked in red, and cAMP phospho-sites, located within predicted NLSs. This was in line with both PKA and CK2 involved in regulating pathways wherein ARID5B is active^30,41^. The nuclear export site is shown as black/yellow stripes. The truncating variants of individuals 16 and 20 are shown too, truncated proteins of 521 and 967 aa long, respectively. (**E**) Overview of mutated ARID5B proteins tested. In brown variants from cohort individuals, in blue a variant from the gnomAD database, in chartreuse green, a variant that was previously associated with ASD^14^, finally, in black are the designed variants. (**F**) Microscope images of the cellular localization of various variants in different growth conditions to assess cellular localization and stability of the phenotype under divergent conditions. Overnight transfected cells were then cultured for 24 hours without serum, after which serum was added 3 hours before staining (+S24h), or no serum was added (SF), or kept in serum-rich medium during the whole experiment (+S). (**G**) Quantification of the expression area of ARID5B variants using a one-way ANOVA (*F*(11, 57) = 20.78, P = 4.37e-16, one-tailed, alpha = 0.05). Post hoc t-tests (one-tailed; increase of surface area for defect nuclear localization) comparing each variant to the control (ARID5B-IsoI) were performed and Bonferroni-corrected. Significant increases were observed for Q522* (*P*-adj. = 1.14 × 10^⁻7^), Q474* (*P*-adj. = 5.68 × 10⁻^5^), Y968* (*p* adj = 0.00125), del1018-1026 (*P*-adj. = 0.036), and G634Afs34 (*P*-adj. = 1.54 × 10^⁻7^). Error bars indicate SD. The grey-blue bar represents wild-type ARID5B, the orange bars are variants identified in our cohort, and the green bars are in-house designed variants. The light yellow bar one of the missense variants found in the genomic studies on ASD. Finally, the red bar represents a truncating gnomAD_V3 variant. *P*-adj. < 0.05 (*), *P*adj. < 0.01 (**), and *P*-adj. < 0.001 (***), indicate adjusted (*P*-adj.) afer post-hoc Bonferroni adjustments. For conciseness of the figure we did not include the exact *P-*value in this figure.

### C-terminal truncating variant shows only a very minor delocalization phenotype

We set up a highly-sensitive, semi-automated blinded quantification of the cellular localization and assessed p.Gln522Ter and the most-C-terminal variant, p.Ala1146CysfsTer15. Whereas p.Gln522Ter showed, as expected, a strong cytosolic delocalization, p.Ala1146CysfsTer15 only showed a very minor but reproducible, cytosolic localization defect (**Fig. 5B**). However, we want to note that the typical localization patterns of p.Ala1146CysfsTer15 is similar to those of wild-type ARID5B, regardless of total protein expressed per cell (**Fig. S2B**), a small portion of cells showed cytosolic localization more similar to p.Gln522Ter (**Fig. S2C**). This could explain the small effect on overall cellular localization of p.Ala1146CysfsTer15 observed in **Figure 5B**.

The long exon 10 harbors 3 ATG sites that have predicted functional Kozak-sequences that have the potential to produce alternative isoforms (**Fig. S3A**). In mouse embryonic brain lysates, proteins of the expected sizes of these alternative exons react with anti-ARID5B antibodies (**Fig. S3B**). In addition, we have found evidence for alternative isoform expression from C-terminally tagged, overexpressed ARID5B (**Fig. S3C**). To investigate if such isoforms would localize in the nucleus, which would suggest the use of a C-terminal nuclear localization signal (NLS), we overexpressed this predicted alternative isoform III and assessed its localization. Indeed, this C-terminal part of ARID5B alone, encoded solely by exon 10, also resided in the nucleus (**Fig. 5C**). A detailed description of potential alternative isoforms can be found in the supplemental information and figure **Fig. S3**.

### Various tools predict a complex nuclear export/import domain in exon 10

To predict protein sequences that could play a role in ARID5B cellular localization, we combined the output of several motif-predicting tools (**Fig. 5D**). NLStradamus *homo Sapiens* predicted two significant nuclear localization (NLS) sites in the N-terminal half of the protein (**Fig. 5D**)^18^. However, a broader almost significant C-terminal domain was indicated by NLS-Mapper, Prosite and MotifScan, with two NLSs flanking a nuclear export signal (NES) in the region covering amino acids 999-1030 of ARID5B isoform-I, potentially forming a complex NLS/NES/NLS (NNN) domain (**Fig. 5D**)^19–21^. Notably, one of the missense variants in our cohort (p.Ser266Asn) resides on the flank of the predicted N-terminal NLS (**Fig. 5D**).

Considering the truncating variants, most truncations lead to the complete loss of the C-terminal predicted NNN, including p.Tyr968Ter, and thus we hypothesized that cellular localization for variants that lost the NNN would be similarly affected (**Fig. 5D**).

### Cohort and designed ARID5B variants suggest a dominant C-terminal nuclear localization domain

We generated nine more ARID5B variants tagged with a FLAG peptide to investigate consequences on cellular localization (**Fig. 3E**). First, we confirmed that all overexpressed truncated variants result in proteins of the expected size (**Fig. S2D**). In addition to p.Gln522Ter and the more C-terminal p.A1146CfsTer15, we also included shorter and intermediate truncated variants from our cohort (c.1423C>T, p.Gln475Ter and c.2904C>A, p.Tyr968Ter) and the NLS-located missense (p.Ser266Asn). Further, we added several designed variants aiming to explore the mechanism of cytosolic localization in more detail, including one with a small deletion (p.del1018-1026) in the middle of the NNN domain and additional missense in the primary NLS signal (p.Ser264/Ser266Asn) that affects a potential regulatory serine in the N-terminal NLS domain near the variant of individual 3. We also added one C-terminal truncating variant from gnomAD V3 (p.Glu634AlafsTer29) and one missense variant derived from the genomic ASD study, p.Ala702Pro^14^. As expected, truncating variants from the cohort (p.Gln474Ter, p.Tyr968Ter) and gnomAD truncating (p.Gly634AlafsTer29) variants that completely lose the NNN domain located in the cytosol (**Fig. 5F**). We further quantified this by measuring the (increased) average surface expression per cell (**Fig. 5G**). Of note, we could not find any changes in expression patterns caused by stress or altered growth conditions following temporarily depleting cells of serum (**Fig. 5E**).

The p.Ser266Asn cohort variant located in the N-terminal NLS did not lead to more FLAG-tagged protein outside the nucleus. Additional mutation of p.Ser264Leu (p.Ser264Leu/Ser266Asn) did not change this finding (**Fig. 5F,G**). Strikingly, a small deletion in the middle of the NNN (p.del1018-1026) had a surprisingly strong effect, completely delocalizing ARID5B in the cytosol (**Fig. 5F,G**). Other designed variants with altered charges of the N-terminal NLS (p.Lys255Asp/Lys256Asp) and one variant with two serines mutated in the first NLS of the NNN domain (p.Ser1002/1015Asp) did not change the cellular localization (**Fig. 5F,G**). Neither did the ASD-associated p.Ala702Pro variant affect the nuclear localization of ARID5B, all having a punctate nuclear signal that is typical for wild type ARID5B (**Fig. S2E**). In summary, our data reveals a conserved C-terminal ARID5B protein domain that can regulate ARID5B cellular localization, explaining cellular delocalization of truncated ARID5B variants.

## Discussion

We have gathered evidence that truncating *ARID5B* variants, especially those that cluster within exon 9 and at the beginning of the exceptionally long last exon 10 of *ARID5B,* escape NMD, cause a neurodevelopmental disorder with GDD, ID and speech delay. Additional features included kidney defects and recurrent infections. Two individuals had episodes of CNS inflammation and two infants had persistent pulmonary hypertension, being the main cause of death of a 6 months old boy. Facial dysmorphic features most frequently include a short, bulbous nose, noted for half of the individuals, often accompanied by a shorter/lower nasal bridge. Several individuals presented with micrognathia, ptosis and some with a prominent forehead.

While none of the variants that cluster in the first quarter of exon 10 were inherited (14/19 were tested), three out of four of the truncating variants outside the cluster were inherited and the fourth one could not be addressed, which suggest that these variants could have unique deleterious effects for some phenotypical aspects, possibly with milder ID. While one mother was not diagnosed, two others were affected.

Truncations that lead to the loss of fewer domains may lead to reduced loss of function compared to variants that loos the complete exon 10 coding coding RNA, particularly if the additional domain(s) are still functional. Since LoF variants that lead to complete NMD are so rare, as discussed above, we opt that the lack of losing a conserved domain of unknown function, spanning amino acid 610 and 650 of ARID5B isoform one, can play a role. This domain may still be intact in all inherited truncating variants, while being lost in variants that cluster at the beginning of exon 10.

Another possible mutation type specific phenotype could be macrocephaly, with which two individuals with missense variants presented but none of the individuals with truncating variants. Also, four out of eight individuals with ASD had variants that did not cluster in exon 10, while three out of four individuals with a missense variant had ASD. Several missense and one LoF of TV1 were identified in individuals with ASD. We did not find clear other patterns in genotype-phenotype relationships.

None of the missense variants were identified in gnomAD V4, and the affected residues are conserved between mouse and human (p.Ile54Met, individual 1), or between zebrafish, mouse and human (p.Arg186Trp, p.Ser266Asn, p.Arg335Ser, indivduals 2-4). All were present in missense intolerant regions, and three were confirmed to be *de novo*, suggesting (neuro)developmental importance. However, p.Ser266Asn can be found substituted into a glycine (12X) in gnomAD V4 and had unique phenotypes (no ASD, mild microcephaly) compared to the other missense variants. The other three individuals with missense variants and ASD, have variants residing in known critical domains like the BAH and ARID domain, and two of them presented with macrocephaly (BAH and ARID domain variants).

*ARID5B* is predicted to be haploinsufficient considering the gnomAD V4 database, having a loss-of-function (LoF) observed/expected upper bound fraction (LOEUF) of 0.21, with the current threshold for V4 being 0.6^11^. However, *ARID5B* variants that are predicted to lead to the complete loss of function (LoF) *ARID5B* via NMD of both *ARID5B* transcript variants (TV1 and TV2) are much more sparse than LoF in general. To our knowledge, out of >800.000 individuals in gnomAD V4, and theoretically dozens more that could have been identified through gene matching, we could only find two individuals (both in gnomAD V4) that carried variants that are predicted to lead to NMD of both *ARID5B* TV1 and TV2. In addition, variants in the exceptionally long final exon, exon 10, generally escape NMD^7^.

Therefore, the translational consequences can be particularly divergent for *ARID5B,* with LoF variants in a large part of the coding regions escape mRNA decay, or otherwise, only affecting TV1, not TV2, leading in both cases to only a partial loss of endogenous ARID5B protein functions and/or abnormal protein variants.

Both complete (TV1+TV2) LoF variants were identified in gnomAD, while in our cohort, and in many large genomic ASD and ID studies, no additional similar variants were identified. This extremely low frequency could emphasize a strong deleterious effect compared to other *ARID5B* variants, perhaps generally too strong to be compatible with survival. The presence of variants in gnomAD is sometimes interpreted as meaning that they are unlikely to cause neurodevelopmental disorders, but at very low frequencies, one should be cautious interpreting gnomAD in such a way^22^–it should be kept in mind that these individuals for instance may have a mosaic variants, or are affected by these variants. We conclude that only complete genotype and phenotype characterization of heterozygous individuals with LoF variants can determine if such ultra-rare variants indeed lead to limited or variably penetrant phenotypes. For now, we expect that complete *ARID5B* LoF (affecting both transcript variants) will generally be as deleterious, or more deleterious as the truncating variants described here.

A few other potentially deleterious *ARID5B* variants have been previously described. One individual with a nonsense variant affecting only TV1 (p.Gln84Ter) presented with a neurodevelopmental disorder, with ID, in combination with seizures, pituitary defects, a severe short stature^23^, which are mostly phenotypes we have not observed with other *ARID5B* variants. Several other variants, both LoF affecting only TV1, and missense variants in the C-terminal half of ARID5B, have been associated with autism spectrum disorder (ASD)^13,13,14^ and several more can be found in gnomAD V4.

Several immune and inflammatory roles have been ascribed to *ARID5B.* For instance, in adaptive natural killer (NK) cells, *ARID5B* mRNA levels correlate with the expression of electron transport chain genes, oxidative metabolism and Interferon gamma levels^24^.ARID5B’s role as an immune mediator has been supported in aged monocytes, in which increased *ARID5B* expression correlates with atherosclerosis. In these aged monocytes, *ARID5B* is upregulated by immune activation and acts upstream as an activator of lipid metabolism and inflammatory genes. Moreover, the knockdown of *ARID5B* also reduced their migration, while increased expression correlated with increased mobility^25,26^. *ARID5B* transcription itself is under the influence of *ARIEL* (*ARID5B*-inducing enhancer-associated long noncoding RNA/*XLOC_005968)* which is also up-regulated in TAL1-positive types of T-ALL and acts as an enhancer RNA by facilitating *ARID5B* enhancer-transcription complexes^27^. A SNP in intron 4 of *ARID5B*, wherein two alternative exons reside^28^, has been linked to autoimmune diseases, specifically rheumatoid arthritis (RA [OMIM: 180300]) and Graves disease (GD [OMIM: 275000])^29^. In our cohort, two individuals had severe neuroinflammatory episodes (individuals 23 and 26) and a third individual (20) showed regressive loss of speech externalization from the age of two and was hospitalized for several weeks due to a prolonged fever with unknown cause around the age of 10. Given the rarity of cerebellitis and encephalomyelitis in pediatrics, the often precarious clinical situation that may arise, and given the numerous associations between *ARID5B* and immune system function, we think that it is critical to emphasize that truncating *ARID5B* variants are likely causally related to the recurrent CNS inflammatory episodes and other immune perturbations experienced by this subset of patients.

Furthermore, we discovered that the long exon 10 ORF has an independent nuclear localization domain that is lost for most variants following truncation, except the two most C-terminal variants. These former variants localize in the cytosol, and not in the nucleus of HEK293T cells, in which ARID5B is normally found with a typical punctate nuclear localization (**Fig 5, Fig. S2**). We observed this effect particularly for variants that have translation termination codons upstream of a predicted, complex nuclear localization sequence (NNN). The NNN is very likely involved in ARID5B nuclear localization, in particular because we could drastically diminish ARID5B nuclear localization by a small, designed deletion in the center of the broad NNN domain. Moreover, the predicted isoform-III, which lacks two predicted NLSs that are present in the former half of the main ARID5B isoform, localized in the nucleus, while the loss of the last 42 amino acids (p.A1146CfsTer15, individual 29) of the C-terminus did not substantially affect cellular localization (**Fig. 5B,F,G**). Still, it remains uncertain if the removal of the center of the NNN is directly involved, or that it affects protein localization indirectly by changing the (local) protein conformation. Furthermore, since serine phosphorylation often plays a role in nuclear localization, for example of IKAROS, an ARID5B coregulator^30,31^, we substituted two serines in the NNN-domain but that did not affect the cellular localization of ARID5B.

Escaping *ARID5B* NMD may affect its function in at least three ways as compared to complete LoF. Firstly, the loss of domains encoded by exon 10 may cause only a partial LoF, with the truncated protein still executing some of its functions. Considering the genotype-phenotype relationship, we would like to emphasize that truncating variants that cluster in the first quarter of exon 10 lose all three distinguishable conserved protein domains (or regions) encoded by exon 10 (**Fig. 3**), while the most C-terminal variants may not lose the function of at least two of these domains. Secondly, the truncated protein may have gained functions harmful to neurodevelopment. Thirdly, transcripts that bypass NMD may still express the alternative isoform(s) encoded by exon 10, and their functions may therefore not be lost, reducing the deleterious effect of LoF variants. Investigating the protein localization of ARID5B, isoform expression and truncated protein expression in other cell types and our mice, would surely help address these questions, but are currently challenged by a lack of antibodies that effectively bind to the N-terminal half of ARID5B.

Three separate mouse models with deletions of exons encoding portions of the BRIGHT domain showed strain-dependent sub-lethality frequencies, growth retardation and were lean^32–34^. *Arid5b* mutant mice only deviate postnatally in weight, as adults when they become smaller than wild-type mice^33^. Both homo- and heterozygous *Arid5b* mutant mice are resistant to high-fat diet-induced weight gains. Such resistance to diet-driven weight gains has also been observed in mice lacking ARID5B solely in fat cells by Fabp4-Cre induced deletion of exon 6^34^, which seems to be the consequence of triglyceride and lipid metabolism being downstream of ARID5B^34,35^. Even though, in humans, a SNP in an ARID5B motif has been associated with obesity, and mice lacking ARID5B are lean (or at least are protected against diet-induced weight gains)^34–36^, we did not observe clear trends in weight or growth defects in our cohort, with the individuals falling within a broad spectrum of weight and height. However, 4 individuals presented with IUGR, while later in life, individuals may catch up or even have a taller than average stature. In our mice, their reduced birth weight is at least partially compensated for into adolescence as well. Heterozygous mice seem to catch up, while homozygous mice are too vulnerable to make it through postnatal phases without support.

This discrepancy can be due to several factors. Firstly, the human SNP may affect the binding of other transcription factors, not only ARID5B function. Furthermore, mice that lack ARID5B completely likely have unique phenotypes that could differ from variants that are truncating and escape NMD and/or have gain-of-functions. Therefore, comparing between truncating variants, NMD-causing ‘gene-trap’ mice, or overexpression models should be done cautiously.

We also found subtle changes in the open field test, with, perhaps unexpectedly, the mice spending more time in the center of the arena in the open field test, a phenotype also observed for other ASD/ID NDD mouse models, for example Fragile X syndrome (*Fmr1* disruption), Bosch-Boonstra-Schaaf Optic Atrophy Syndrome (*Nr2f1* disruption), *AUTS2*-related syndrome (*Auts2* disruption)^10,16,17^, and in other mouse models that have established cognitive limitations^37^. An overall lack of awareness, eventual in combination with a lower overall drive to explore could prompt spending more time in the center of an open field, likely even in the absence of an anxiolytic phenotype. Further characterization of our newly generated Arid5b*^emQ522*^* mice, and comparison with gene-trap models can reveal truncation specific deregulated mechanisms that could further pinpoint disease mechanisms specific for truncating variants that locate at the beginning of exon 10, like Arid5b*^emQ522*^*. Challenging the immune system of these mice is a logical future approach to assess their immunological vulnerabilities.

ARID5B also plays a role during chondrogenesis by promoting the production of collagen type II in a manner dependent on SOX9 and involving the activation of SIRT1^38^. This suggests that ARID5B contributes to extracellular matrix (ECM) formation and may support the maintenance of tissue integrity. Although homozygous mice are smaller than their heterozygous counterparts, subtle alterations in ECM formation or phenotypes manifesting only under specific challenges remain to be investigated. Such mechanisms could help explain the variably expressed systemic manifestations—such as hydronephrosis or immunological abnormalities—which may arise not only from direct immune cell perturbations but also indirectly through impaired barrier function and altered cell adhesion secondary to collagen dysregulation.

In conclusion, we have shown that truncating variants, including frameshift variants in exon 9, leading to stop-gains that cluster in the first quarter of the disproportionally long exon 10 of *ARID5B*, cause a neurodevelopment syndrome with a spectrum of phenotypes, including kidney issues, hypotonia and feeding problems, most frequently mild ID and with DD, speech and language delay/disorder and behavioral issues. A broad or bulbous nose, often with a short and low nasal bridge was observed. Other distinctive features included ptosis, radial hand deviations and foot/toe abnormalities. There is also a possible susceptibility to episodes of recurrent infections, often without a known cause, of the urinary tract, upper airways and inner ear, and two rare cases of CNS neuroinflammation, with one individual poorly responding to therapy being treated palliatively at the time of writing. Two infants presented with persistent pulmonary hypertension, leading to the death of individual 19 at six months of age. Truncating mutations escape NMD and showed cellular localization defects that we attribute to the loss or impairment of a C-terminal nuclear localization signal/nuclear export domain. Mice with truncating mutations in *ARID5B* show developmental and behavioral abnormalities. Missense variants in critical domains like the BRIGHT domain and truncating variants in the distal part of exon 10 very likely affect neurodevelopment, but their consequences may require individual assessment due to variable deleteriousness and partially unique phenotypes.

## Author Contributions

H.v.H wrote the manuscript and designed the experiments. H.v.H, N.R, A.G, A.P, and J.C. performed the experiments. H.v.H., N.R. and A.P. analyzed the experiments, P.C. & H.v.H. supervised the project, H.v.H. and P.C. recruited the cohort, and K.B. edited the manuscript. All other authors contributed by reviewing the manuscript and providing clinical information and/or exome analyses.

## Acknowledgements And Funding

Part of this work was performed under the Care4Rare Canada Consortium funded by Genome Canada and the Ontario Genomics Institute (OGI-147), the Canadian Institutes of Health Research, Ontario Research Fund, Genome Alberta, Genome British Columbia, Genome Quebec, and Children’s Hospital of Eastern Ontario Foundation.

Genetic testing of individuals with the c.1419del (p.Asn434LysfsTer45) and the c.1365dupA (p.Glu456ArgfsTer31) was performed within the framework of the GAD (*Génétique des Anomalies du Développement*) collection and approved by the appropriate institutional review board of Dijon University Hospital (DC2011-1332). The work was supported by grants from Dijon University Hospital, the ISITE-BFC (PIA ANR), the European Union through the FEDER programs (PERSONALISE), and the Burgundy-Franche-Compté regional council (INTEGRA). The sequencing platform at the CNRGH was supported by the France Génomique National infrastructure, funded as part of the “Investissements d’Avenir” program, managed by the Agence Nationale pour la Recherche (contract ANR-10-INBS-09). The whole genome sequencing performed at the CNRGH was funded by the Laboratory of Excellence GENMED (Medical Genomics) Grant No. ANR-10-LABX-0013, managed by the National Research Agency (ANR) as part of the Investment for the Future program.

The CRISPR-Cas9 images (thumbnail) (image “Cas9 gene”) was reproduced from the original work by the author (2024) under the Creative Commons Attribution-ShareAlike 4.0 International License. No modifications were made. The inner ear illustration (thumbnail) was provided by MedicalGraphics.de and is used under a Creative Commons Attribution-NoDerivatives 4.0 International License (CC BY-ND 4.0). No modifications were made to the original image. The drawing of the lungs Medical illustration (thumbnail) by Patrick J. Lynch and C. Carl Jaffe, Yale University School of Medicine, licensed under CC BY 2.5.

Megaureter illustration courtesy (thumbnail) of the Children’s Hospital of Philadelphia, used under public domain. The image “Cartoon brain illustration” (thumbnail) was obtained from the public domain collection on Rawpixeland is free to use without restriction under the CC0 Public Domain Dedication.

## Ethics Declaration

The study was conducted in accordance with the Declaration of Helsinki and approved by the Ethics Committee of CHU Sainte-Justine (protocol code 015-853:4072 and date of approval 26 May 2015). Written informed consent from the participants’ legal guardian/next of kin was provided in accordance with the national legislation and the institutional requirements for the provided images to be published and donated blood to be used for scientific purposes.

## Data And Code Availability

The published article includes all datasets generated or analyzed during this study. Raw imaging data, constructs, and plasmid maps will be provided upon request. R code for statistics in figures 4C, 4G, and 5G, is available at https://github.com/Hjvanheesbeen/ARID5B_Cohort_Statistics.

---

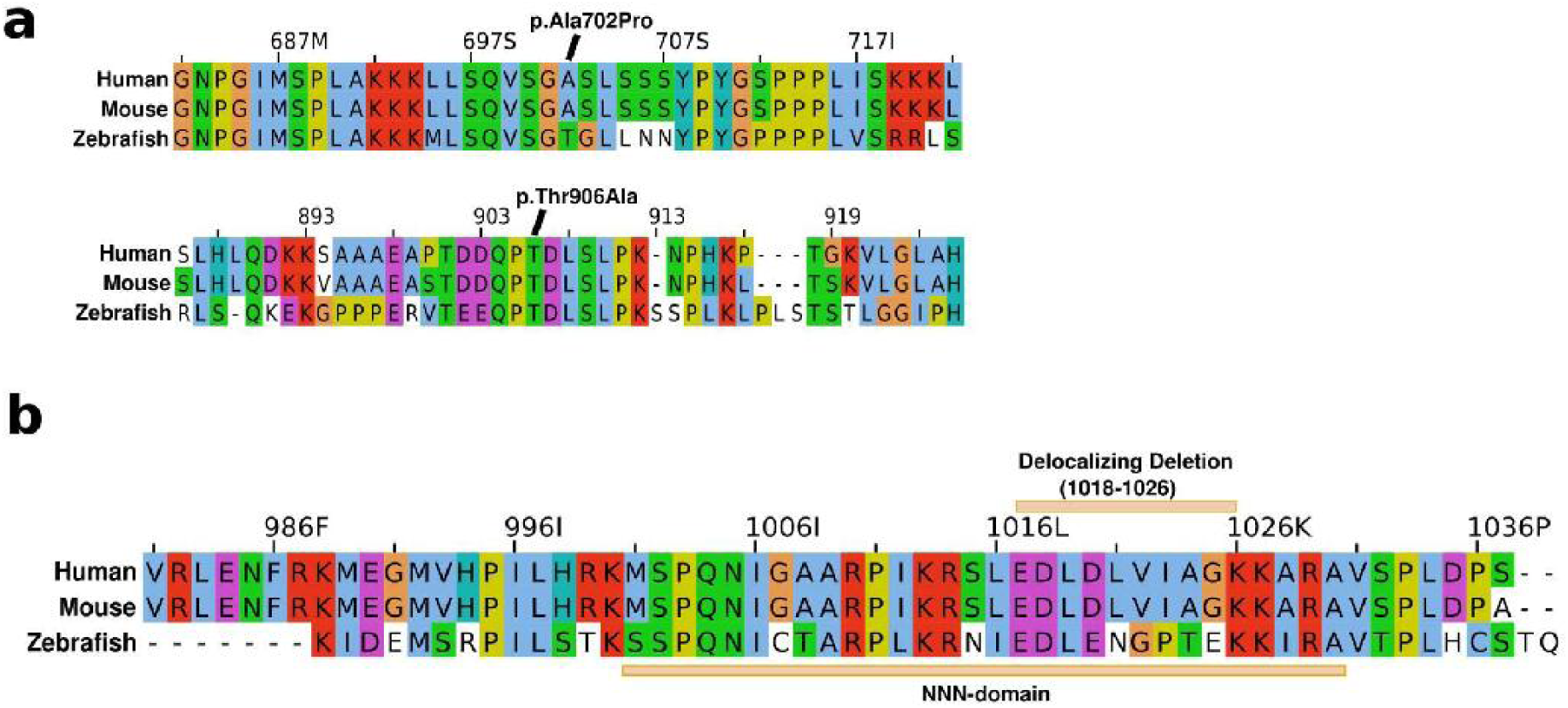

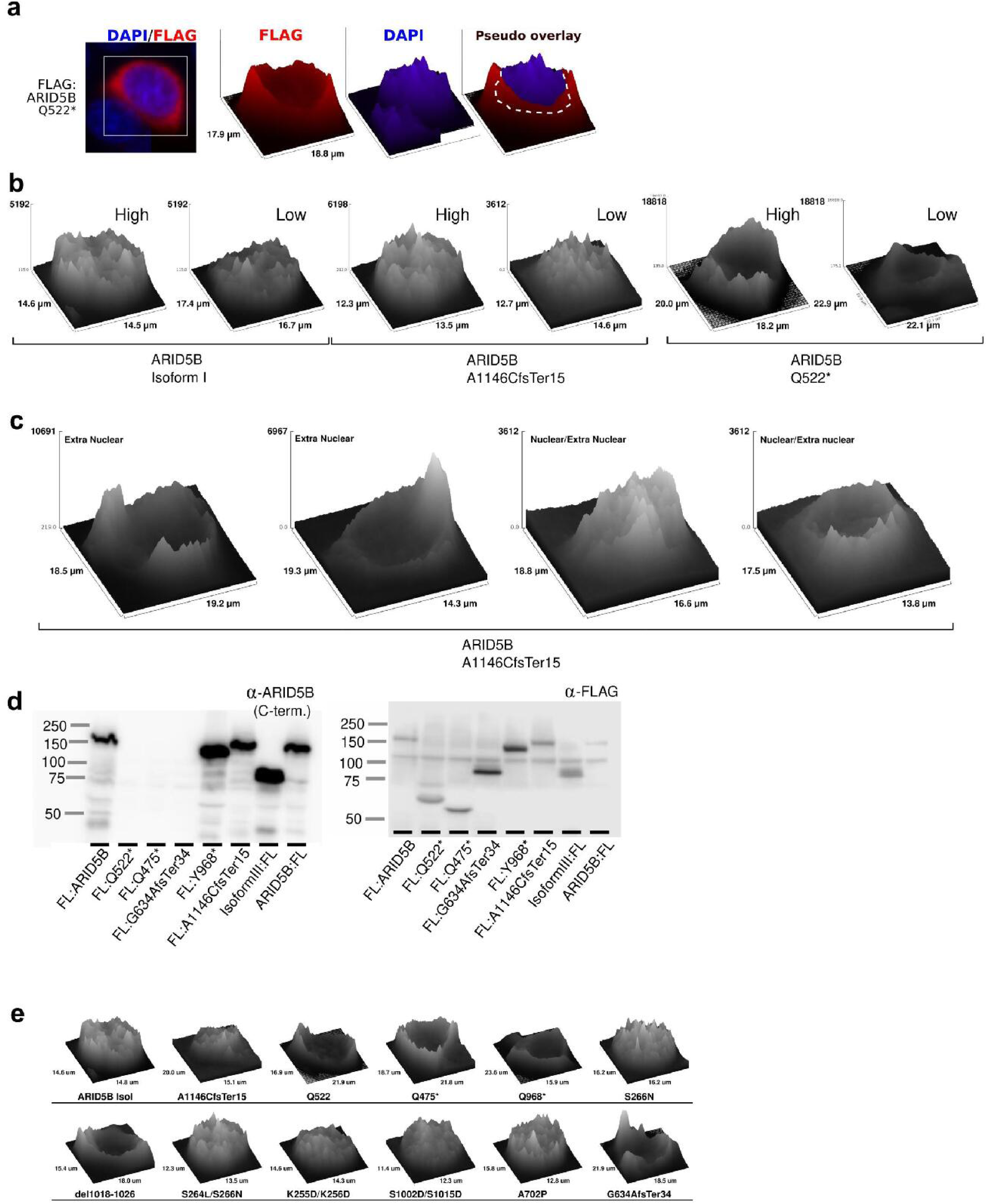

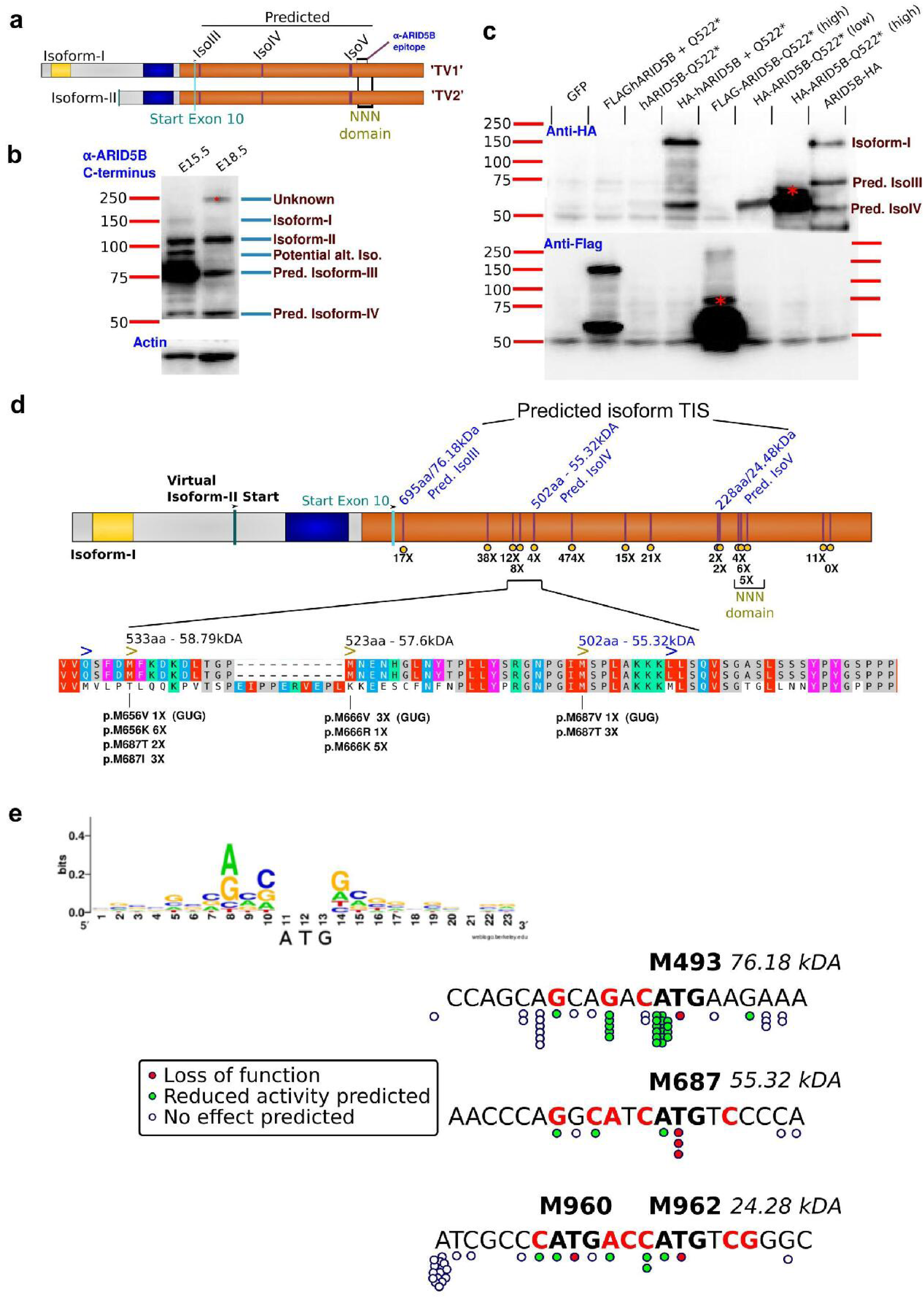

## References

1. Deák G, Cook AG. Missense Variants Reveal Functional Insights Into the Human ARID Family of Gene Regulators. Journal of Molecular Biology. 2022;434(9):167529. doi:10.1016/J.JMB.2022.167529

2. Kosho T, Okamoto N, Collaborators CSSI. Genotype-phenotype correlation of Coffin-Siris syndrome caused by mutations in SMARCB1, SMARCA4, SMARCE1, and ARID1A. American Journal of Medical Genetics Part C: Seminars in Medical Genetics. 2014;166(3):262–275. doi:10.1002/ajmg.c.31407

3. Vals MA, Õiglane-Shlik E, Nõukas M, et al. Coffin–Siris Syndrome with obesity, macrocephaly, hepatomegaly and hyperinsulinism caused by a mutation in the ARID1B gene. Eur J Hum Genet. 2014;22(11):1327–1329. doi:10.1038/ejhg.2014.25

4. Schrier Vergano SA. ARID2, a milder cause of Coffin-Siris Syndrome? Broadening the phenotype with 17 additional individuals. American Journal of Medical Genetics Part A. 2024;194(6):e63540. doi:10.1002/ajmg.a.63540

5. Lelieveld SH, Reijnders MRF, Pfundt R, et al. Meta-analysis of 2,104 trios provides support for 10 new genes for intellectual disability. Nature Neuroscience 2016 19:9. 2016;19(9):1194–1196. doi:10.1038/nn.4352

6. Gudlaugsdottir S, Boswell DR, Wood GR, Ma J. Exon size distribution and the origin of introns. Genetica. 2007;131(3):299–306. doi:10.1007/S10709-007-9139-4/TABLES/3

7. Lindeboom RGH, Supek F, Lehner B. The rules and impact of nonsense-mediated mRNA decay in human cancers. Nature genetics. 2016;48(10):1112. doi:10.1038/NG.3664

8. Philippakis AA, Azzariti DR, Beltran S, et al. The Matchmaker Exchange: A Platform for Rare Disease Gene Discovery. Human mutation. 2015;36(10):915. doi:10.1002/HUMU.22858

9. Schindelin J, Arganda-Carreras I, Frise E, et al. Fiji: an open-source platform for biologi-cal-image analysis. Nature Methods 2012 9:7. 2012;9(7):676–682. doi:10.1038/nmeth.2019

10. Sobreira N, Schiettecatte F, Valle D, Hamosh A. GeneMatcher: A Matching Tool for Connecting Investigators with an Interest in the Same Gene. Human Mutation. 2015;36(10):928–930. doi:10.1002/HUMU.22844

11. Karczewski KJ, Francioli LC, Tiao G, et al. The mutational constraint spectrum quantified from variation in 141,456 humans. Nature 2020 581:7809. 2020;581(7809):434–443. doi:10.1038/s41586-020-2308-7

12. von Ehr J, Oberstrass L, Yazgan E, et al. Arid5a uses disordered extensions of its core ARID domain for distinct DNA- and RNA-recognition and gene regulation. Journal of Biological Chemistry. 2024;300(7):107457. doi:10.1016/j.jbc.2024.107457

13. Satterstrom FK, Kosmicki JA, Wang J, et al. Large-Scale Exome Sequencing Study Implicates Both Developmental and Functional Changes in the Neurobiology of Autism. Cell. 2020;180(3):568. doi:10.1016/J.CELL.2019.12.036

14. Krumm N, Turner TN, Baker C, et al. Excess of rare, inherited truncating mutations in autism. Nat Genet. 2015;47(6):582–588. doi:10.1038/ng.3303

15. Hori K, Nagai T, Shan W, et al. Heterozygous Disruption of Autism susceptibility candidate 2 Causes Impaired Emotional Control and Cognitive Memory. PLOS ONE. 2015;10(12):e0145979. doi:10.1371/journal.pone.0145979

16. Saré RM, Levine M, Smith CB. Behavioral Phenotype of Fmr1 Knock-Out Mice during Active Phase in an Altered Light/Dark Cycle. eNeuro. 2016;3(2):ENEURO.0035-16.2016. doi:10.1523/ENEURO.0035-16.2016

17. Contesse T, Ayrault M, Mantegazza M, Studer M, Deschaux O. Hyperactive and anxio-lytic-like behaviors result from loss of COUP-TFI/Nr2f1 in the mouse cortex. *Genes*, Brain and Behavior. 2019;18(7):e12556. doi:10.1111/gbb.12556

18. Nguyen Ba AN, Pogoutse A, Provart N, Moses AM. NLStradamus: a simple Hidden Markov Model for nuclear localization signal prediction. BMC Bioinformatics. 2009;10:202. doi:10.1186/1471-2105-10-202

19. Sigrist CJA, De Castro E, Cerutti L, et al. New and continuing developments at PROSITE. Nucleic acids research. 2013;41(Database issue). doi:10.1093/NAR/GKS1067

20. Kosugi S, Hasebe M, Tomita M, Yanagawa H. Systematic identification of cell cycle-dependent yeast nucleocytoplasmic shuttling proteins by prediction of composite motifs. Proceedings of the National Academy of Sciences of the United States of America. 2009;106(25):10171–10176. doi:10.1073/PNAS.0900604106/SUPPL_FILE/0900604106SI.PDF

21. Pagni M, Ioannidis V, Cerutti L, et al. MyHits: improvements to an interactive resource for analyzing protein sequences. Nucleic acids research. 2007;35(Web Server issue). doi:10.1093/NAR/GKM352

22. Gudmundsson S, Singer-Berk M, Watts NA, et al. Variant interpretation using population databases: Lessons from gnomAD. Human Mutation. 2022;43(8):1012–1030. doi:10.1002/humu.24309

23. Martinez-Mayer J, Vishnopolska S, Perticarari C, et al. Exome Sequencing Has a High Diagnostic Rate in Sporadic Congenital Hypopituitarism and Reveals Novel Candidate Genes. The Journal of Clinical Endocrinology & Metabolism. 2024;109(12):3196–3210. doi:10.1210/clinem/dgae320

24. Cichocki F, Wu CY, Zhang B, et al. ARID5B regulates metabolic programming in human adaptive NK cells. The Journal of Experimental Medicine. 2018;215(9):2379. doi:10.1084/JEM.20172168

25. Liu D, Zhang XX, Li MC, et al. C/EBPβ enhances platinum resistance of ovarian cancer cells by reprogramming H3K79 methylation. Nature Communications. 2018;9(1):1739. doi:10.1038/s41467-018-03590-5

26. Saare M, Tserel L, Haljasmägi L, et al. Monocytes present age-related changes in phospholipid concentration and decreased energy metabolism. Aging cell. 2020;19(4). doi:10.1111/ACEL.13127

27. Tan SH, Leong WZ, Ngoc PCT, et al. The enhancer RNA ARIEL activates the oncogenic transcriptional program in T-cell acute lymphoblastic leukemia. Blood. 2019;134(3):239. doi:10.1182/BLOOD.2018874503

28. Kent WJ, Sugnet CW, Furey TS, et al. The Human Genome Browser at UCSC. Genome Research. 2002;12(6):996–1006. doi:10.1101/GR.229102

29. Okada Y, Terao C, Ikari K, et al. Meta-analysis identifies nine new loci associated with rheumatoid arthritis in the Japanese population. Nature Genetics. 2012;44(5):511–516. doi:10.1038/ng.2231

30. Ge Z, Han Q, Gu Y, et al. Aberrant ARID5B expression and its association with Ikaros dysfunction in acute lymphoblastic leukemia. Oncogenesis 2018 7:11. 2018;7(11):1–10. doi:10.1038/s41389-018-0095-x

31. Uckun FM, Ma H, Zhang J, et al. Serine phosphorylation by SYK is critical for nuclear localization and transcription factor function of Ikaros. Proc Natl Acad Sci U S A. 2012;109(44):18072–18077. doi:10.1073/pnas.1209828109

32. Lahoud MH, Ristevski S, Venter DJ, et al. Gene Targeting of Desrt, a Novel ARID Class DNA-Binding Protein, Causes Growth Retardation and Abnormal Development of Re-productive Organs. Genome Research. 2001;11(8):1327–1334. doi:10.1101/GR.168801

33. Whitson RH, Tsark W, Huang TH, Itakura K. Neonatal mortality and leanness in mice lacking the ARID transcription factor Mrf-2. Biochemical and Biophysical Research Communications. 2003;312(4):997–1004. doi:10.1016/j.bbrc.2003.11.026

34. Whitson RH, Li SL, Zhang G, Larson GP, Itakura K. Mice with Fabp4-Cre ablation of Arid5b are resistant to diet-induced obesity and hepatic steatosis. Molecular and cellular endocrinology. 2021;528. doi:10.1016/J.MCE.2021.111246

35. Yamakawa T, Sugimoto K, Whitson RH, Itakura K. Modulator recognition factor-2 regu-lates triglyceride metabolism in adipocytes. Biochemical and Biophysical Research Communications. 2010;391(1):277–281. doi:10.1016/j.bbrc.2009.11.049

36. Claussnitzer M, Dankel SN, Kim KH, et al. FTO Obesity Variant Circuitry and Adipo-cyte Browning in Humans. The New England journal of medicine. 2015;373(10):895. doi:10.1056/NEJMOA1502214

37. Usui N, Berto S, Konishi A, et al. Zbtb16 regulates social cognitive behaviors and neo-cortical development. Transl Psychiatry. 2021;11:242. doi:10.1038/s41398-021-01358-y

38. Hata K, Takashima R, Amano K, et al. ARTICLE Arid5b facilitates chondrogenesis by recruiting the histone demethylase Phf2 to Sox9-regulated genes. Published online 2013. doi:10.1038/ncomms3850

39. Colin E, Duffourd Y, Tisserant E, et al. OMIXCARE: OMICS technologies solved about 33% of the patients with heterogeneous rare neuro-developmental disorders and negative exome sequencing results and identified 13% additional candidate variants. Frontiers in Cell and Developmental Biology. 2022;10:1021785. doi:10.3389/FCELL.2022.1021785

40. Silk M, Petrovski S, Ascher DB. MTR-Viewer: identifying regions within genes under purifying selection. Nucleic Acids Research. 2019;47(W1):W121–W126. doi:10.1093/NAR/GKZ457

41. Baba A, Ohtake F, Okuno Y, Yokota, et al. PKA-dependent regulation of the histone ly-sine demethylase complex PHF2-ARID5B. Nature cell biology. 2011;13(6):668–675. doi:10.1038/NCB2228

